# Allele specific expression and gene regulation explain transgressive thermal tolerance in non-native hybrids of the endangered California tiger salamander (*Ambystoma californiense*)

**DOI:** 10.1101/772020

**Authors:** Robert D. Cooper, H. Bradley Shaffer

## Abstract

Hybridization between native and non-native species is an ongoing global conservation threat. Hybrids that exhibit traits and tolerances that surpass parental values are of particular concern, given their ability to outcompete the native parent. It is crucial to understand the mechanisms that drive these transgressive hybrid traits to diagnose and develop strategies to manage hybrid populations. Here, we explore several aspects of the hybridization between the endangered California tiger salamander (*Ambystoma californiense*; CTS) and the introduced barred tiger salamander (*Ambystoma mavortium*; BTS). We assayed critical thermal maximum (CTMax) to compare the ability of CTS, BTS and hybrids to tolerate acute thermal stress, and found that hybrids exhibit a wide range of CTMax values, with 40% (6/15) able to tolerate temperatures greater than either parent. We quantified the genomic response of each individual to discover and compare thermal abatement strategies. We found that CTS and BTS have strikingly different numbers and tissue-specific patterns of overall gene expression, with hybrids expressing intermediate values. We evaluated transgressive and variable phenotypes by uncovering regulatory mechanisms that give rise to these unique traits. F1 hybrids display abundant and variable degrees of allele specific expression (ASE), likely arising from extensive compensatory evolution in gene regulatory mechanisms of the parental lineages. We found that the proportion of genes with allelic imbalance in individual hybrids correlates with their CTMax, suggesting that BTS-biased expression confers improved thermal tolerance. We discuss the implications of these findings with respect to ongoing management of CTS in the face of future climate change.

## Introduction

### Threat of Invasive Species and Hybridization

The introduction and establishment of invasive species represents one of the most significant threats to vertebrate biodiversity worldwide (Bellard, Cassey, & Blackburn, 2016). Non-native species pose immediate threats due to competition or predation of native flora and fauna (Gurevitch & Padilla, 2004). However, these non-native species pose a more devastating and much less understood threat when they hybridize with native taxa. Globalization and increased human movement have brought allopatric species into contact that have not evolved isolating mechanisms that reduce or eliminate hybridizing (Mallet, 2005). Hybridization between divergent species is often halted at the F1 stage due to genomic or behavioral incompatibilities. However, instances of hybridization that do produce viable offspring often results in genomic introgression. This invasion of non-native genes into local populations typically results in the decline of the native parents as they are replaced demographically by hybrid offspring (Rhymer & Simberloff, 1996). The most common outcome of non-native introgression is genetic swamping, in which native local genomes are replaced by those of mixed ancestry. This process often culminates in genomic extinction, where no pure native genotypes persist (Allendorf, Leary, Spruell, & Wenburg, 2001; Ellstrand & Rieseberg, 2016). A recent review of hybridization across multiple taxa concluded that 69 out of 143 studies reported hybridization to be an extinction threat (Todesco et al., 2016).

Genetic swamping can be intensified by anthropogenically-induced environmental perturbations including climate change (Gómez, González-Megías, Lorite, Abdelaziz, & Perfectti, 2015). Organisms that are successful invaders often have intrinsic abilities that allow them to rapidly adapt to novel environments. Alien species may also bring genes that are specifically adapted to local landscapes where they are introduced, resulting in non-native admixture fortifying hybrid animals against future environmental change (Hamilton & Miller, 2016; Kovach, Luikart, Lowe, Boyer, & Muhlfeld, 2016; Pfennig Karin S., Kelly Audrey L., & Pierce Amanda A., 2016; Rieseberg, Archer, & Wayne, 1999). This threat can be particularly severe when hybrids possess traits or tolerances that surpass that of their parental species.

### Transgressive Hybrid Phenotypes

Heterosis and transgressive segregation are two well-documented phenomena that can produce hybrids with phenotypic trait values that exceed either parent. Heterosis typically involves first generation hybrid crosses (F1) and is thought to result from dominant or overdominant alleles that mask recessive alleles from the alternative parental line, thus producing exaggerated phenotypes (Comings & MacMurray, 2000; Dobzhansky, 1952). Transgressive segregation occurs at, or after, the F2 stage (second generation hybrid crosses), when meiosis can produce recombinant genotypes. This phenomenon occurs more frequently than previously assumed; 97% of plant and 78% of animal hybrid studies report transgressive phenotypes (Rieseberg et al., 1999). Dittrich-Reed and Fitzpatrick (2013) presented a comprehensive genetic model that can explain the occurrence and persistence of these transgressive hybrid phenotypes. However, these models do not explicitly evaluate mechanisms that may favor F1 hybrids before they progress to the F2 stage. Selection occurring at this initial F1 stage may be critical in understanding the rapid success, or demise, of hybrid populations in nature.

Though often viewed independently, heterosis and transgressive segregation are linked processes that can produce transgressive hybrid phenotypes. Both phenomena are predicted to produce more exaggerated hybrid phenotypes with increasingly divergent parental lineages (Guindon, Martin, Cravero, & Cointry, 2019). Furthermore, heterosis has been used in many studies to predict the successful establishment of transgressive phenotypes in subsequent generations (Chahota, Kishore, Dhiman, Sharma, & Sharma, 2007; Guindon et al., 2019; Khan, Mahbub, Reza, Shirazy, & Mahmud, 2016; Kumar, Jeberson, Singh, & Sharma, 2017). These studies of agricultural crops highlight the importance of examining trait values in the F1 hybrid stage to make meaningful predictions about long-term success. In the case of wild non-native hybrids, it is similarly useful to view phenotypic traits in F1 hybrids to understand potential advantages and adaptations that may arise through heterosis and/or transgressive segregation. These traits may experience positive selection, promoting the success of F1 individuals and thus facilitate further hybrid crosses with a greater potential of establishing permanent transgressive hybrid populations. In this study we utilize F1 hybrid phenotypes to understand the forces promoting hybridization between an endangered salamander and an introduced congener in central California.

### Hybridization in the California Tiger Salamander System

The California tiger salamander (“CTS”, *Ambystoma californiense*) is endemic to the grassland ecosystems of central California and is protected under both federal and state law (U.S. Fish and Wildlife Service, 2004). One of the primary threats to CTS is the rapid introgression of non-native alleles from the introduced barred tiger salamander (*Ambystoma mavortium*; “BTS”). BTS were intentionally introduced into the Salinas Valley (Monterey County, California, USA) in the 1950’s and 1960’s for the fishing bait industry (Riley, Shaffer, Voss, & Fitzpatrick, 2003). Hybridization between CTS and BTS occurs readily, and these hybrids are fertile, allowing introgression past the F1 stage (J. R. Johnson, Fitzpatrick, & Shaffer, 2010). These non-native genotypes reach high frequencies in native populations more rapidly than is expected based on models of neutral diffusion (Fitzpatrick & Shaffer, 2007b), and ongoing landscape genomic analyses confirm that the hybrids are expanding and are present in all three of the CTS distinct population segments (McCartney-Melstad and Shaffer, unpublished data). Several studies indicate the presence of transgressive phenotypes in these hybrid salamanders, which may promote their success in the wild. Hybrids were found to have increased larval survival compared to CTS and BTS, affording them a direct fitness advantage (Fitzpatrick & Shaffer, 2007a), and Searcy et al. (2016) concluded that hybrids do not serve the same ecological function as native salamanders, resulting in less biodiverse breeding ponds. This pattern is likely driven by the enhanced growth rate of hybrid larvae, which enables them to prey on aquatic organisms typically too large for native CTS (Ryan, Johnson, & Fitzpatrick, 2009; Searcy et al. 2016). This rapid growth rate also results in larger hybrid salamanders at metamorphosis, which has been shown to significantly improve lifetime fitness in CTS (J. R. Johnson, Ryan, Micheletti, & Shaffer, 2013; Searcy, Gray, Trenham, & Shaffer, 2014). F1 hybrids also have superior locomotor capabilities compared to either CTS or BTS (J. R. Johnson, Johnson, & Shaffer, 2010). These studies suggest that CTS x BTS hybrids interact with environmental variation in novel ways, likely facilitated by transgressive phenotypes.

### Thermal Stress and CTS

In this study we focus on temperature stress given its critical role in tiger salamander survival and its potential to promote salamanders with enhanced thermal tolerance in the wild. Semi-arid grasslands are a challenging environment for pond-breeding amphibians to carry out their entire life-cycle, and are therefore rarely inhabited by salamanders. Sparse rains and high temperatures are strong selective pressures on amphibians, and members of the tiger salamander complex have evolved adaptations to these conditions which enable them to occupy habitat that precludes virtually all other salamanders in North America (Petranka, 1998; Stebbins, 2003). For CTS in particular, limited annual precipitation leads to shallow breeding ponds that experience rapid, large temperature fluctuations, subjecting metamorph and aquatic larvae to extreme heat (Holland, Hayes, & McMillan, 1990; Pounds, 2001). After metamorphosis, emerging CTS take temporary refuge in nearby rodent burrows while they wait for the necessary rainfall to continue their upland migration (Loredo, Van Vuren, & Morrison, 1996). These initial opportunistic refugia must be adequate to survive up to six months of extreme Central Valley heat since inadequate burrow selection may expose salamanders to lethal temperatures (Searcy and Shaffer; in Review). CTS must then select permanent burrows for summer aestivation to survive extreme summer temperatures (Trenham, Bradley Shaffer, Koenig, & Stromberg, 2000). It is therefore likely that hybrid tiger salamanders with transgressive thermal tolerance may enjoy increased fitness in this challenging landscape. We investigate the role of acute thermal stress on CTS, BTS and hybrid salamanders using a critical thermal maximum (CTMax) assay. CTMax is defined as the upper thermal limit of a species temperature tolerance, above which the organism is unable to function. Detecting differences in this trait allows us to compare the efficacy of each group’s thermal tolerance mechanism.

### Methods to Investigate Transgressive Phenotypes

In this study we outline a research program that can be used to identify transgressive hybrid phenotypes and the genetic mechanisms that produce them. First, we quantify differences in physiological response to environmental stimuli between hybrids and the parental taxa under common environmental conditions. Specifically we investigate the effects of acute thermal stress on CTS, BTS and hybrid crosses to identify potential differences in thermal tolerance. Second, we examine the genomic response to specific environmental conditions in parental and hybrid groups. Differential Gene Expression (DGE) enables us to compare the expression profiles of CTS, BTS and hybrids in response to thermal stress. Finally, we examine regulation of gene expression to better understand transgressive or variable phenotypes in F1 hybrids. Gene regulation is a complex process involving any number of promotors located on the same DNA molecule near the gene of interest (*cis-*) and regulatory elements located elsewhere that are not physically linked to the focal gene (*trans-*). Analysis of gene expression in the parental species and of parent-specific gene copies in the F1 hybrids allows us to identify genes that are governed by *cis-*, *trans-*, or any combination of the two regulatory factors. We employ these methods to assign regulatory modes to each gene of interest in hybrid tiger salamanders, then make inferences about the evolutionary history and mechanisms that give rise to transgressive hybrid phenotypes. From these three levels of analysis we seek to answer the following questions: 1.) At the whole organism physiological level, are there differences in thermal tolerance between CTS, BTS and hybrids? 2.) At the underlying genetic level, are there differences in gene expression between CTS, BTS and F1 hybrids exposed to near-lethal thermal stress? 3.) Can we identify transgressive or variable phenotypes in the F1 hybrids? 4.) If we find transgressive or variable phenotypes in F1 hybrids, what genomic mechanisms drive their extreme phenotypes?

## Materials and Methods

### Study Population

We used tiger salamanders from a captive research colony housed at the University of California, Los Angeles. These individuals were either wild-caught or captive bred and were housed individually in a climate controlled room for at least 4 years ensuring that individuals were sexually mature and fully acclimated to the laboratory thermal regime. Salamanders were fed pinky mice and/or crickets twice per week, and housed following approved UCLA animal care protocols (ARC #2013-011-11).

For each salamander in the study we measured: Snout to vent length (SVL), total length, tail length, tail height, jaw width, and mass. Body condition was calculated as the residuals of the SVL and mass linear model. Sex was determined at the time of dissection based on presence of ovarian follicles (Winne & Ryan, 2001). Salamanders of each genotype were randomly assigned to experimental (heated) and control groups. This resulted in 10 CTS (5 control, 5 heat), 12 BTS (6 control, 6 heat), 25 F1-hybrids (13 control, 12 heat), and 4 Backcross/F2-hybrids (1 control, 3 heat) for the CTMax experiment. Despite the limited sample size, backcross/F2-hybrids were included in the physiological experiment to compare CTMax in later generation crosses with that of F1 hybrids. These individuals were not included in downstream expression analyses.

### Physiology and CTMax

Critical Thermal Maximum (CTMax) was determined for each salamander in the experimental treatment. This metric has been used in many physiological experiments on salamanders to describe a species’ tolerance of near-lethal temperatures (Burke & Pough, 1976; Spotila, 1972). CTMax has been shown to correlate well with other measures of thermal tolerance, and is a useful tool in predicting population persistence in response to changing climate (Araújo et al., 2013; Huey et al., 2012). We assessed CTMax using the loss of righting response (LRR), a technique that has been used extensively in ectotherms (Delmas, Baudry, Girondot, & Prevot-Julliard, 2007; Gvozdík, 2011; Sanabria, Quiroga, & Martino, 2012) including CTS (J.R. Johnson, Johnson, Shaffer, 2010). The inability to right itself represents a loss of function that prevents the organism from escaping stressful conditions or fleeing from a predator, therefore LRR is a whole-animal performance measure with direct ecological significance (W. I. Lutterschmidt & Hutchison, 1997).

To assay CTMax, individuals were placed on a moist paper towel substrate in an opaque plastic container under a 100w ceramic infra-red heat emitting bulb (Zoo Med brand) (Layne & Claussen, 1982; Young & Gifford, 2013). Salamanders were constantly misted during the heating trials to avoid desiccation and were monitored for signs of behavioral abnormalities. Temperatures were constantly monitored with an infra-red temperature gun (Amprobe IR-720) positioned 3 inches from the dorsum. After dorsal temperatures reached 30°C individuals were assayed for loss of righting response (LRR). LRR was achieved when an individual was unable to right itself after 30 seconds on its back on a moist surface (W. I. Lutterschmidt & Hutchison, 1997; Young & Gifford, 2013). At this time temperature was recorded with an IR temperature gun 1cm from the dorsal and ventral surface of the body. Control individuals were placed in the same plastic containers in the same room as the CTMax trials and flipped on their back to parallel the LRR assay, but were kept at constant room temperature of 21°C.

Previous work has suggested that measurements taken using non-contact infrared thermometers accurately estimate core body temperature in amphibians, without the need of invasive cloacal probing (Rowley & Alford, 2007; Tracy, Christian, & Tracy, 2010), though some inconsistencies have been detected in reptiles (Carretero, 2012). We therefore calibrated our temperature measurements using museum specimens to accurately estimate internal core temperature from dorsal and ventral surface measurements. We used eight ethanol-preserved specimens of CTS, BTS and hybrids within the size range of our experimental animals as thermal models. Temperatures were recorded using IR temperature guns positioned 1cm from the dorsal and ventral sides of each salamander. A thin thermocouple probe was also inserted into the center of the body cavity to record true internal temperature. We heated each specimen for one hour and recorded temperatures every 3 minutes during this period. We then created linear models that estimate the internal body temperature as a function of dorsal and ventral skin temperatures using the software R (v3.3.2 CRAN).

We analyzed CTMax to address our primary question: do hybrid salamanders possess transgressive thermal tolerance abilities compared to both parental lineages? To test this, we first compared the CTMax of CTS and BTS. If no difference was detected, we pooled CTS and BTS into a parental group and compared the CTMax of the parents with that of the hybrids. We use this approach to identify differences in both the mean and variance of CTMax.

### RNA Lab Protocol

Immediately after CTMax was achieved, or after 60 minutes for controls, individuals were anesthetized in a 5% Benzocaine/ water solution for 5 minutes, decapitated to ensure instant euthanasia, and brain and muscle tissues were harvested. Entire brains and muscle tissue from the rear left leg were taken and immediately placed in RNALater (Ambion) solution and stored at - 20°C until RNA extraction.

Tissues were homogenized using a bead shaker, and then extracted using a modified Purelink RNA Mini Kit-TRIzol spin column protocol. Samples were individually marked with dual index iTru barcodes to allow for multiplexing (Glenn et al., 2016). Libraries were prepared using Kapa mRNA Stranded Library Prep kits and standard protocols. Libraries were pooled and split across six lanes of 100bp single-end reads on an Illumina HiSeq4000 at the QB3 Genomics Sequencing Laboratory in Berkeley, CA.

### Trimming and Mapping

Raw sequences were trimmed for adapter sequence and low quality bases using Trimmomatic (Bolger, Lohse, & Usadel, 2014). Quality and repetitive regions were assessed across all samples using FASTQC (Andrews, 2010). Reads were mapped to the recently published Mexican Axolotl (*Ambystoma mexicanum*) genome (Nowoshilow et al., 2018), a close phylogenetic relative of CTS and BTS (Shaffer & McKnight, 1996, Shaffer and McCartney-Melstad, unpublished). We mapped reads to this genome using STAR (Dobin et al., 2013) which is designed to map transcripts to a genome by efficiently accounting for gap junctions created by intronic sequences. Mapped reads were counted for differential expression analysis using HTSeq-count with exons as the target feature (Anders, Pyl, & Huber, 2015). Reads that mapped to multiple exons were discarded as ambiguous.

### Differential Expression

Differential expression (DE) was calculated for muscle and brain tissue separately using the R package DESeq2 (Love, Huber, & Anders, 2014). A recent comparison demonstrated fewer false discoveries, and more consistent identification of DE genes with DESeq2 compared to similar software (Seyednasrollah, Laiho, & Elo, 2015). We used the exon count output from HTseq since DESeq2 has an internal method for normalizing expression data based on gene-wide dispersion (Love et al., 2014). To reduce the number of genes observed, we removed transcripts with mean count less than 10 from the analysis. We also removed backcross/F2 hybrid individuals from subsequent expression analyses given the limited sample size of these groups. DESeq2 was then run with a model that included one factor (∼groups) with six levels. Each level consisted of unique genotype and treatment combinations (CTS.heat, CTS.control, BTS.heat, BTS.control, F1.heat, F1.control). This allowed for independent contrasts between the various levels, allowing us to determine genes differentially expressed in response to temperature within each genotype class (CTS, BTS, F1). We also tested the overall treatment effect by calculating the average expression of the three genotypes in response to temperature.

### Allele Specific Expression

We analyzed allele specific expression (ASE) to identify differences in BTS / CTS allelic expression in F1 hybrids. To quantify ASE we first called variant sites following the Genome Analysis ToolKit (GATK) best practices pipeline (Auwera et al., 2013). We removed duplicated reads using picardtools, then used the mapped reads to call individual genotypes which were then compared across samples to find polymorphic SNPs. We filtered variant sites using VCFtools to find alleles that were fixed in the pure genotype samples (fixed different in CTS and BTS), filtering out all other variants. We then kept only SNPs that were also heterozygous in all F1 individuals. This enabled us to create a reference file which contained a conservative list of loci that are present in F1s and diagnostic between CTS and BTS. We then filtered the VCF SNP database to only include these loci and used the GATK tool ASEReadCounter to count the number of copies of reference and alternate alleles in each F1 hybrid (Auwera et al., 2013). We assigned parental origin to each allele using the diagnostic BED file using custom scripts in the software R (v3.3.2 CRAN). We have a reduced risk of reference mapping bias on ASE since both parental genotypes contain a combination of reference and non-reference alleles (Stevenson, Coolon, & Wittkopp, 2013).

### GeneiASE

We used the software package geneiASE to identify SNPs and genes that exhibited significant allele specific expression (Edsgärd et al., 2016). This software uses a beta-binomial distribution to compare the counts of reference and alternate SNPs at heterozygous loci against the null hypothesis that alternate allele counts (altCounts) = reference allele counts (refCounts). The software computes a test statistic (s) which is the log(odds-ratio) of the counts of diagnostic alleles. This test statistic is then summarized over all SNPs within a given feature (i.e. annotated gene) using Stouffer’s method to determine feature wide allelic imbalance. This gene-wide allelic imbalance is then compared to a null distribution generated by taking *k*-samples from the beta-binomial distribution and determining the likelihood of the observed imbalance. This yields a probability that the observed allelic ratios for a given gene are equal, and that observed differences in allelic contribution originated by chance, resulting in a list of genes that exhibit significant bias regardless of experimental treatment (“static ASE”). GeneiASE was also used to identify condition-dependent ASE by comparing F1 allelic bias in the heat treatment to the median allelic counts of the control group. All p-values were adjusted using the Benjamini-Hochberg false discovery correction (FDR ≤ 0.001; Benjamini & Hochberg, 2000). Although geneiASE provides a robust framework for determining genes with significant ASE, it does not report the direction of the bias. We summed the number of CTS and BTS counts for each gene and each sample identified by geneiASE, and took the log_2_ ratio of (BTS / CTS) to determine direction and magnitude of ASE.

### Regulatory Mechanisms

We identified the regulatory mechanisms underlying allelic imbalance in F1 hybrids by comparing the relative expression of parental genotypes to their relative expression within the hybrid individuals (ASE of parental copies). We corrected for differences in library sequence depth by calculating Counts Per Million (CPM) for each gene and for each sample. To identify genes with significant *trans-* regulatory factors, we followed the protocol of McManus *et al*. (2010) by comparing ASE ratios in the F1 hybrids to the ratio of BTS/CTS expression in the parental genotypes using Fisher’s Exact test followed by Benjamini-Hochberg correction for multiple tests (Benjamini & Hochberg, 2000). We then attributed the following regulatory mechanisms to each gene: “conserved“, “*cis* only“, “*trans* only“, “compensatory“ or a combination of “*cis* and *trans*“ regulation (see Figure 1) following the method of McManus *et al*. (2010). However, we determined genes that were differentially expressed between CTS and BTS using the output of DESeq2 as described above. We also determined genes with unbalanced expression of parental alleles using the output from the static geneiASE analysis described above. We believe that these software packages offer a more robust determination of significant genes given our sample design, which incorporates information on individual variation in expression where McManus et al. (2010) used pooled libraries of multiple individuals. We further divided the category of “*cis* and *trans“* into four groups where *cis-* and *trans-* functioned in opposite or similar directions, and where either *cis-* or *trans-* had a greater effect on expression (see Figure 1) following the protocol of Goncalves et al. (2012). All analyses were performed using custom scripts in the software R (v3.3.2 CRAN).

**Figure 1.**
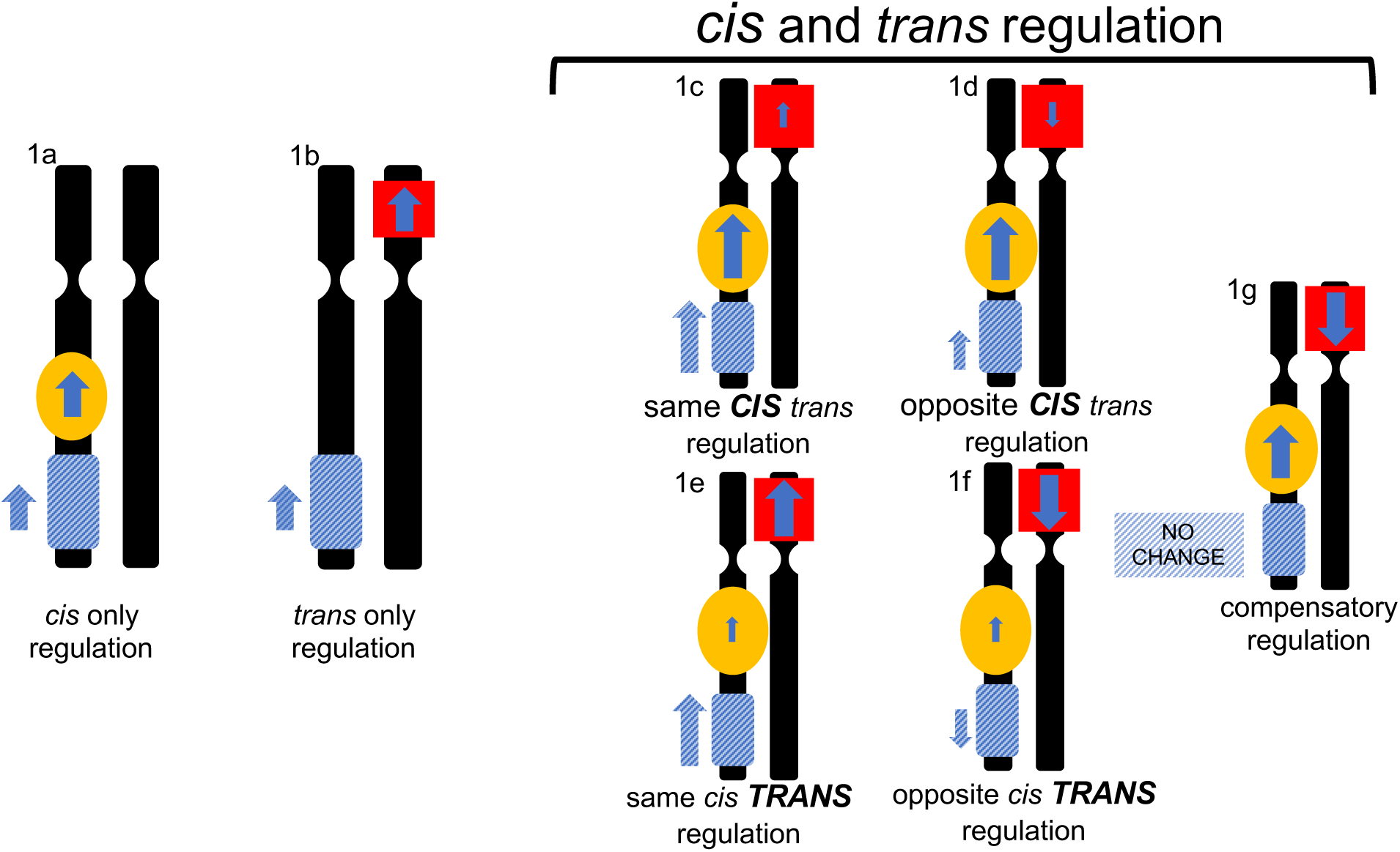
Schematic illustrating the different regulatory modes that were assigned to each gene. Blue hatched box represents a focal gene and the matching blue arrow represents the overall expression of that gene. Yellow ovals are *cis-* regulatory factors which are proximal to the gene of interest. Red squares represent *trans-* regulatory factors that occur elsewhere in the genome, typically unlinked to the focal gene. The “*cis* and *trans* regulation” group has both *cis-* and *trans-* factors acting together in either the same or opposite directions (depicted by the direction of arrows) and with different relative magnitudes (relative size of the arrows, **BOLD** text). The overall effect on expression is represented by the blue hatched arrow next to the focal gene.

## Results

### Physiology and CTMax

Our use of preserved specimens for internal body temperature calibration demonstrated that linear models containing both dorsal and ventral temperatures were highly significant (p < 2.2e-16, df = 139, adj R^2^ = 0.91), and that the removal of either dorsal or ventral temperature produced inferior models (dAIC = 52.28, 23.21, respectively). We therefore calculated internal body temperatures for each individual using the equation: Internal.Temp = -5.16 + 0.37*Dorsal.Temp + 0.75*Ventral.Temp.

In the temperature stress experiment, we detected no difference in CTMax (Welch t-test: t = 0.22, df = 9.00, p = 0.832; Figure 2) between CTS (33.77°C ± 0.39; mean ± SE, n = 5) and BTS (33.64°C ± 0.43, n = 6). Hybrids (34.24°C ± 0.53, n = 15) had a greater mean CTMax than the pooled parental group (33.70°C ± 0.28, n = 11; Welch t-test: t = 2.56, df = 20.66, p = 0.018; Figure 2). F1 hybrids (35.15°C ± 0.61, n = 12) also had a greater mean CTMax then the pooled parental group (Welch t-test: t = 2.16, df = 15.4, p = 0.046) when only F1 hybrids were analyzed, and backcross/F2 hybrids were removed.

**Figure 2.**
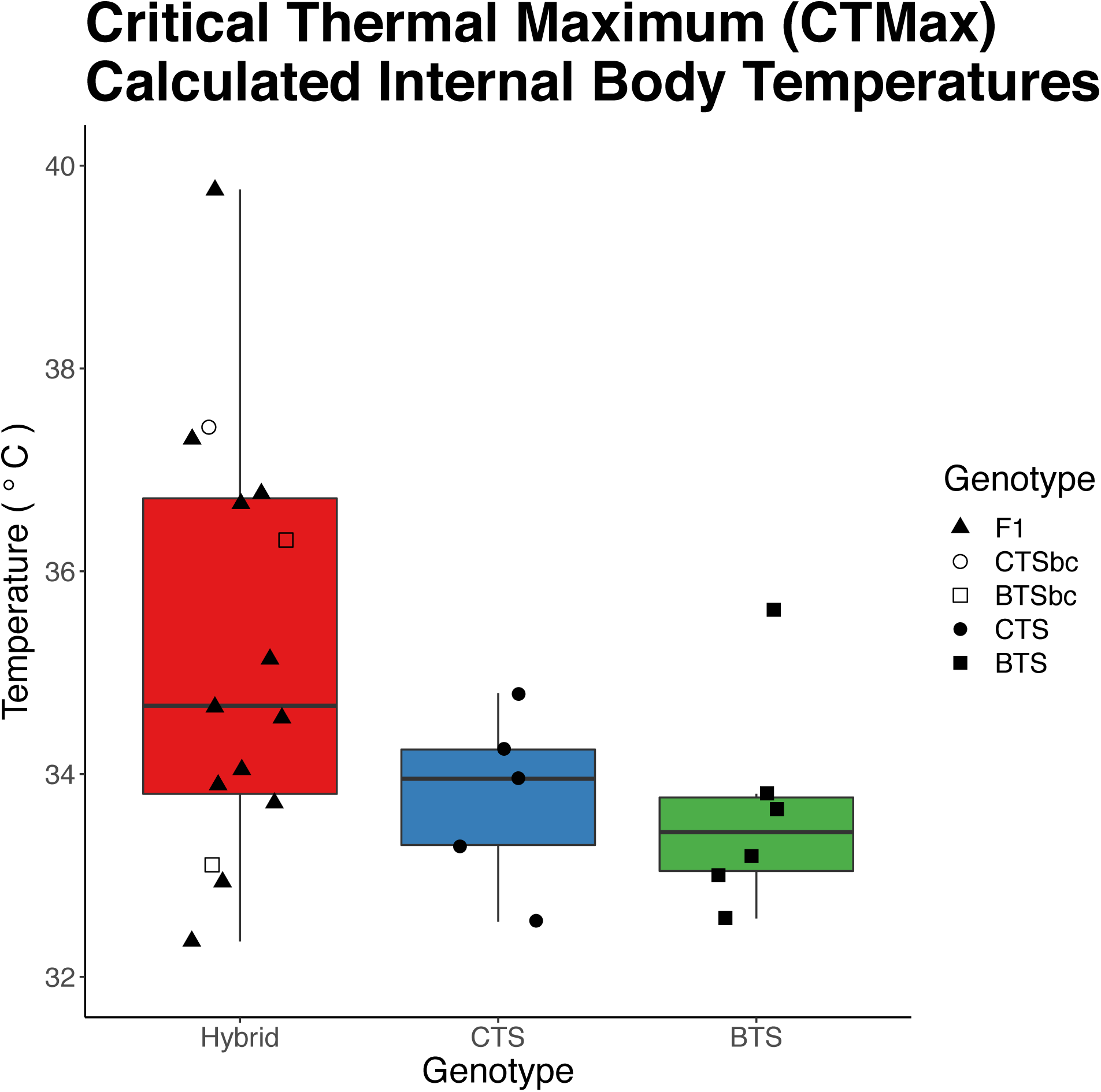
Values of Critical Thermal Maxima (CTMax) by salamander genotype. Barplots display the mean (black horizontal line) and interquartile range (colored rectangle) for each group. Internal body temperatures were calculated using a linear model including dorsal and ventral skin temperatures. CTS is pure California tiger salamander, BTS is pure barred tiger salamander. Hybrid includes three genetic groups: 13 F1 (first generation) hybrids, 2 BTSbc (F1 x BTS offspring) and 1 CTSbc (F1 x CTS offspring).

There was also no significant difference in the variance of CTMax between CTS (sd = 0.87, n = 5) and BTS (sd =1.1, n = 6; F-test: F = 0.67, df = 4, p = 0.721). Hybrids (sd = 2.1, n = 15) did have greater variance in CTMax compared to the combined parental group (F-test: F = 4.88, p = 0.008). When backcross hybrids were removed, F1 hybrids still had greater variance than the parental group (sd = 0.93, n = 11; F-test: F = 5.11, df = 11, p = 0.016). Six of the 15 hybrids (4 F1’s, 1 CTSbc and 1 BTSbc) displayed a greater thermal tolerance than the highest recorded value in either CTS (N = 5) or BTS (N = 6). The mean CTMax of these six transgressive hybrids was 1.77°C greater than the highest recorded CTMax for CTS and BTS combined (35.61°C, BTS). Despite the increased variance, the lower end of the CTMax range did not appear to differ between groups.

### Differential Expression

Differential expression analysis using DESeq2 in the muscle tissue revealed many genes that responded to acute heat stress. Pooling all genotypes (overall treatment effect) revealed 177 DE genes; 106 were up regulated and 71 down regulated. In CTS there were 359 DE genes, 245 up and 114 down. In BTS there 16 DE genes, 8 up and 8 down. In F1 hybrids there were 79 DE genes, 50 up and 29 down regulated (Figure 3).

**Figure 3.**
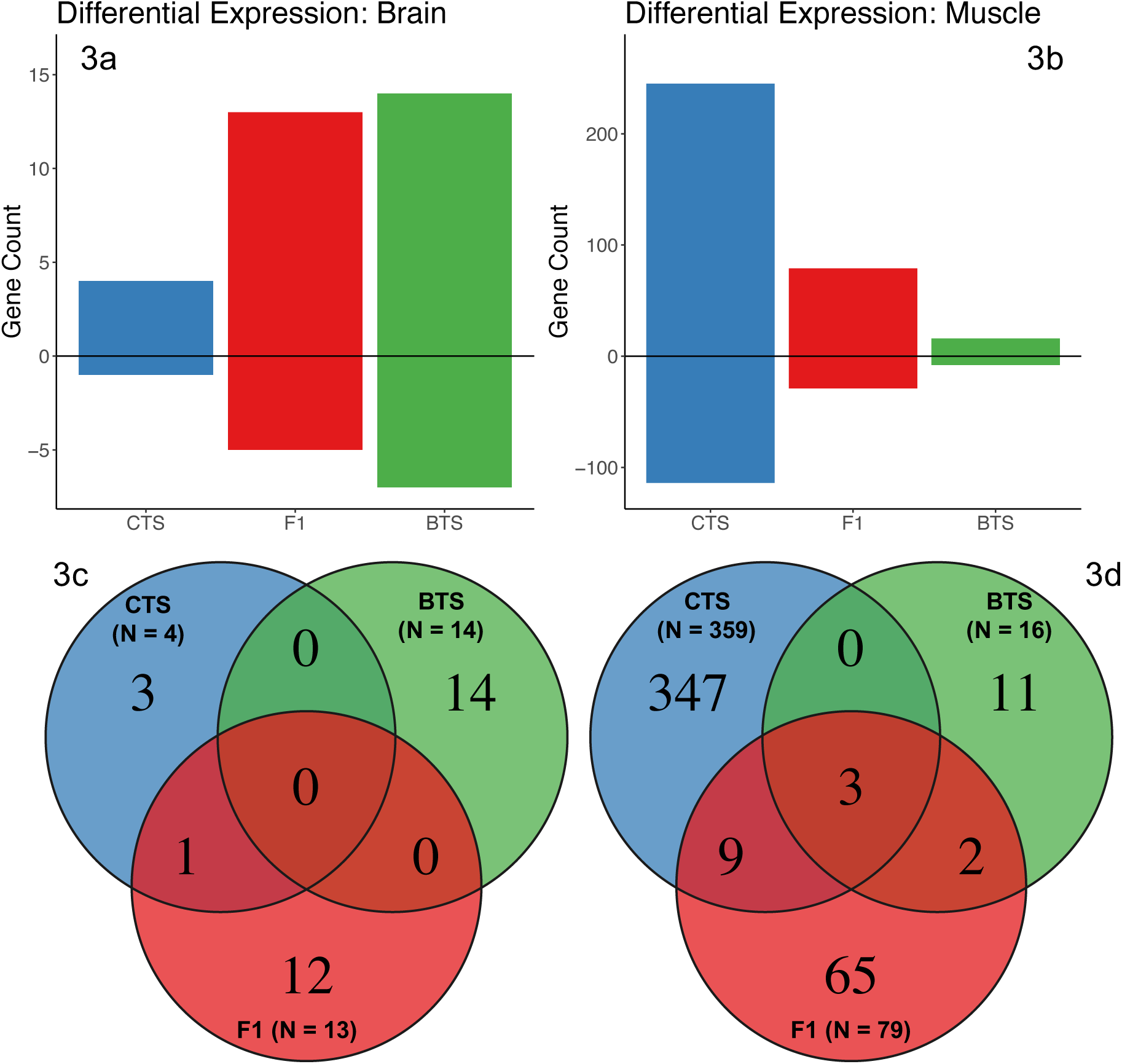
Number of genes differentially expressed in response to acute thermal stress. Figures 3a and 3b depict the number of genes up (positive) and down (negative) regulated in brain (3a) and muscle tissue (3b) for each genotype. Figures 3c (brain) and 3d (muscle) depict the number of genes that were differentially expressed (DE) in response to temperature stress. Values included in overlapping circles indicate genes that were DE in all overlapping groups.

Differential expression in the brain tissue revealed fewer significant genes. Pooling across genotypes (overall treatment effect) yields 32 DE genes, 24 up and 8 down regulated. In CTS there were 4 DE genes, 3 up and 1 down. In BTS 14 genes were DE, 7 up and 7 down. In hybrids a total of 13 genes were DE, 8 up and 5 down (Figure 3).

### Allele Specific Expression

GeneiASE revealed many genes with biased parental expression patterns in F1 hybrids. The median percentages of alleles with significant ASE were 30.7% and 15.7% for muscle and brain tissue respectively. Overall, genes were biased for BTS parental copies in both the brain (log_2_(BTS/CTS) = 0.22; Figure 4a) and the muscle (log_2_(BTS/CTS) = 0.31; Figure 4b), representing a 17% and 25% increase in the expression of BTS gene copies. There was a greater number of genes with significant ASE in the muscle than the brain (ANOVA, df = 45, F = 278.6, p < 2.2e^-16^), and the magnitude of ASE was also greater in the muscle than the brain (Paired t-test by gene, df = 849, p = 5.0e^-5^).

**Figure 4.**
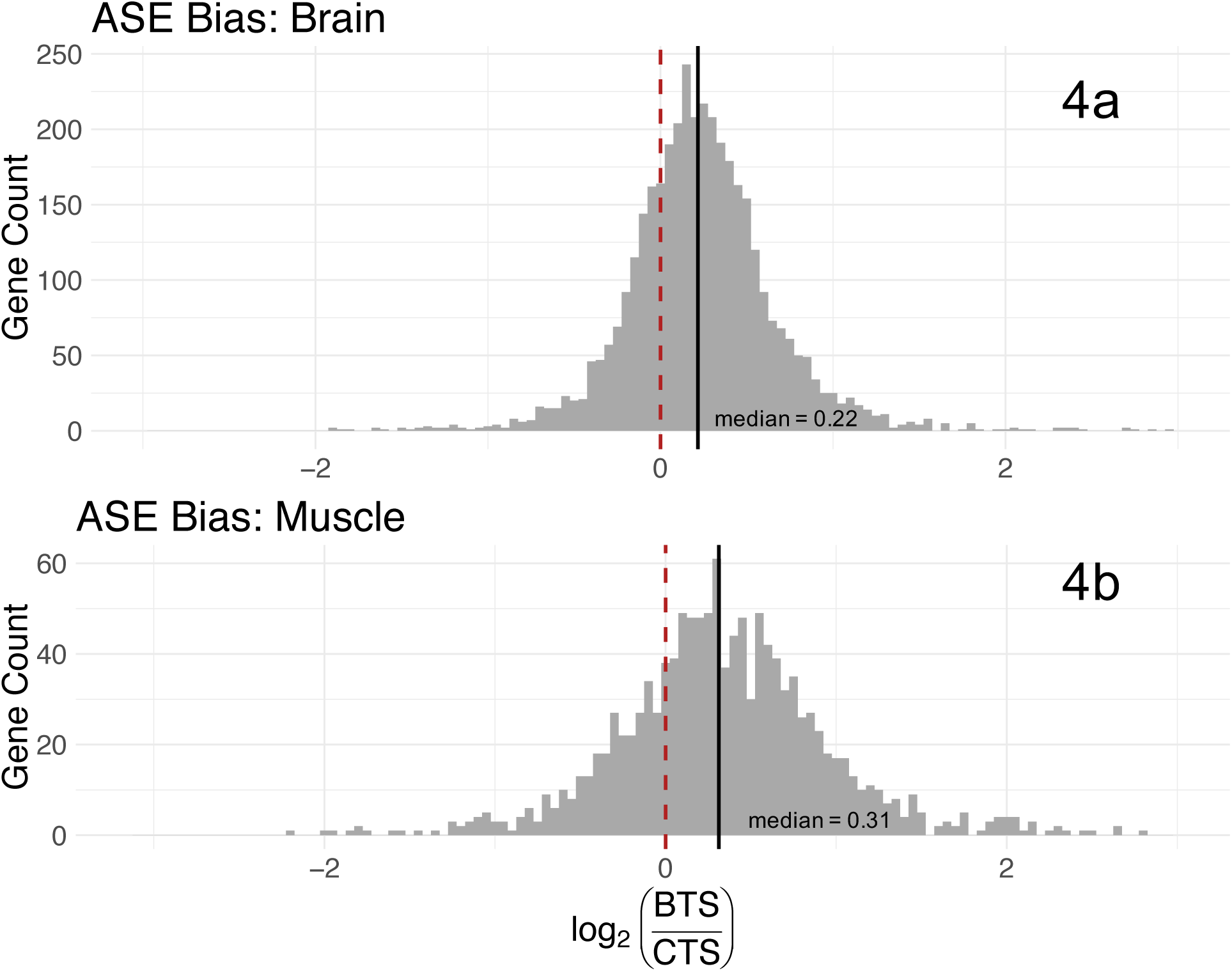
Histograms of the direction and magnitude of allele specific expression (ASE) across all loci that are diagnostic between BTS and CTS. Median log_2_ ratios of BTS/CTS allelic expression were taken for each gene in all F1 individuals. Red dashed line at zero represents equal expression of BTS and CTS alleles. Black line depicts the observed median of all ASE bias across genes for brain (4a) and muscle (4b).

There was no difference in the percentage of genes with significant ASE between hybrids in heat and control treatments in the muscle (ANOVA, df = 23, F = 0.156, p = 0.70) or the brain (ANOVA, df = 20, F = 0.08, p = 0.79). The magnitude of BTS biased ASE was greater in the control hybrids (log_2_ (BTS/CTS) = 0.34) than in the heat treatment (log_2_ (BTS/CTS) = 0.29) in muscle tissue (paired t-test by gene: df = 1240, p = 0.02). The opposite pattern was true in the brain where BTS biased expression was greater in the heat stressed hybrids (log_2_ (BTS/CTS) = 0.23) than in the control individuals (log_2_ (BTS/CTS) = 0.22) (paired t-test by gene, df = 3763, p = 0.04).

There was no significant relationship between percent ASE and CTMax in the muscle (linear regression, F = 0.22, df = 10, p = 0.647). However, there was a strong correlation between the percent of genes with significant ASE and CTMax in the brain (Figure 5a; linear regression: df = 7, F = 13.36, p = 0.036), which explains a significant portion of the variance (R^2^= 0.413, N = 9). There was no significant difference in the magnitude of ASE bias and CTMax in muscle (linear regression: df = 10, p = 0.856) or brain (linear regression: df = 8, p = 0.467) tissue.

**Figure 5.**
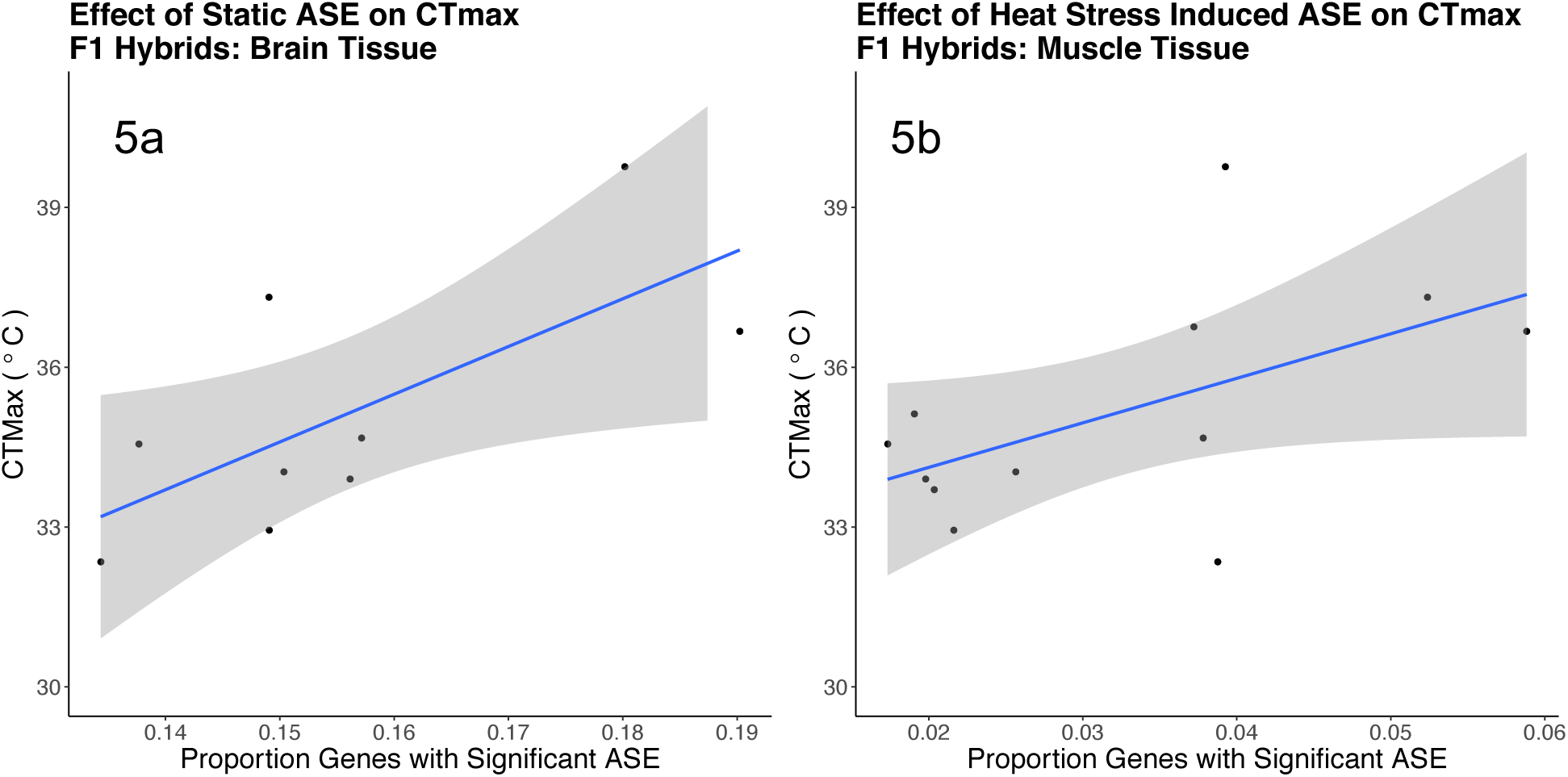
Correlation between each F1 individual’s proportion of genes with significant allele specific expression (ASE) and thermal tolerance. Figure 5a shows the positive trend between an F1 individuals degree of overall ASE bias in brain tissue and its ability to tolerate acute thermal stress. Figure 5b shows a similar trend (although marginally non-significant) between F1 individuals degree of condition-dependent ASE and thermal tolerance in muscle tissue.

The median percentage of genes that exhibited an increase in ASE with response to the heat treatment (condition-dependent) were 3.1% and 0.4% for muscle and brain tissue respectively. There was significantly more condition-dependent ASE in muscle than brain (ANOVA: F = 32.15, df = 18, p = 2.2e^-5^). There was no significant relationship between percent condition-dependent ASE and CTMax in brain (linear regression: F = 1.70, df = 6, p = 0.240), however there is potentially a relationship in the muscle, though the test was marginally non-significant (df = 10, F = 4.29, p = 0.065; Figure 5b).

### Regulatory Mechanisms

Analysis of regulatory mechanisms in F1 hybrids revealed tissue specific patterns. Gene expression in muscle tissue had greater *trans-* only regulation (25.7%) than *cis-* only (19.4%). Muscle tissue also exhibited a great deal of *cis-* and *trans-* factors (54.9%) which were dominated by compensatory mutations (28.8%). Of the genes with both *cis-* and *trans-* regulation, most functioned in opposing directions (18.9%) rather than complimentary directions (7.2%). Brain tissue was more regulated by *cis-* only factors (15.2%) than by *trans-* only elements (4.0%). Additionally, there was a large number of genes with a combination of *cis-* and *trans-* factors (80.8%), again dominated by compensatory regulation (68.6%). In brain tissue, genes regulated by both *cis-* and *trans-* elements predominantly functioned in opposing directions (9.2%) compared with those acting in the same direction (3.0%). All values are presented in Table 1 and displayed in Figure 6.

**Figure 6.**
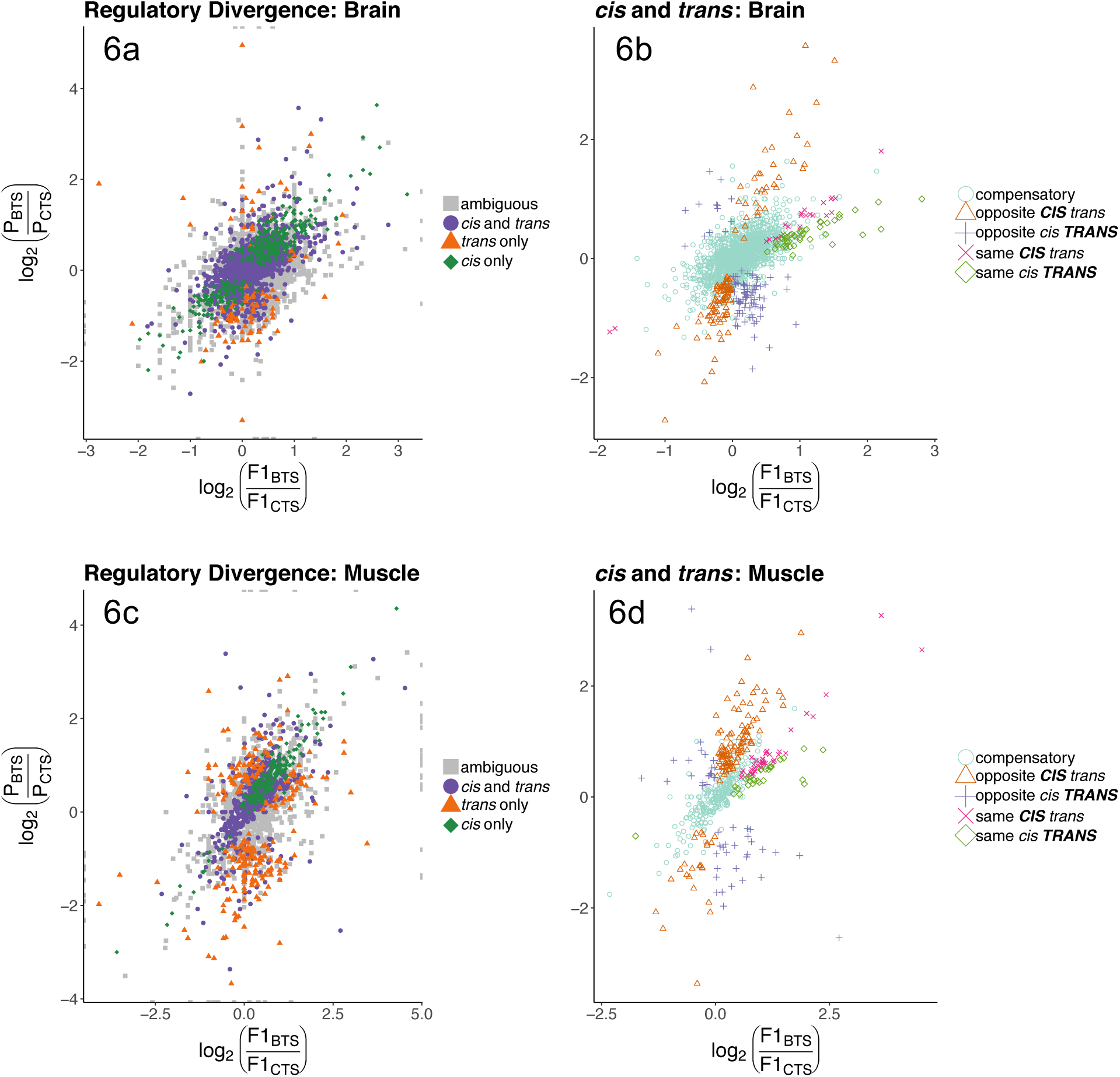
Scatter plots showing the relative expression of CTS and BTS genes in pure parental species and of parent specific gene copies in F1 hybrids. Vertical axis is the log_2_ ratio of BTS/CTS expression of a specific gene averaged across pure parental individuals. Horizontal axis is the log_2_ ratio of BTS/CTS derived gene copies when simultaneously expressed in F1 hybrids. Comparison of these two expression ratios enable us to assign modes of regulation to each gene. Figures 6a and 6c identify genes that are governed by *cis-*, *trans-* or a combination of the two (“*cis* and *trans*”). Figures 6b and 6d expand on the category “*cis* and *trans*”, identifying genes that are more influenced by *cis-* or *trans*-regulation (indicated in **BOLD**), and genes where *cis-* and *trans*-regulation act in the “same” or “opposite” directions. See Figure 1 for further description of the regulatory modes.

**Table 1.** Counts and percentage of genes that demonstrate significant regulatory divergence between BTS and CTS. Genes with significant differential regulation are broken down into three groups: genes with only *cis-*, only *trans-* and genes with a combination of *cis-* and *trans-* regulation. This group “*cis* and *trans*” is further divided by the direction and magnitude of the change in expression. Regulatory divergence may operate in the “opposite” or “same” direction, and the mechanism with the greatest effect on expression is displayed in **BOLD**. See Figure 1 for further explanation of these categories.

Regulatory Mechanisms
|  | Brain |  | Muscle |  |
| --- | --- | --- | --- | --- |
|  | Counts | Percent | Counts | Percent |
| All Genes Considered | 15,899 | 100.0% | 13,945 | 100.0% |
| Conserved | 11,543 | 72.6% | 11,783 | 84.5% |
| Differential Regulation | 2,267 | 14.3% | 931 | 6.7% |
| Ambiguous | 2,079 | 13.1% | 1,231 | 8.8% |
| Genes with Differential Regulation | 2,267 | 100.0% | 931 | 100.0% |
| *cis* only | 344 | 15.2% | 181 | 19.4% |
| *trans* only | 91 | 4.0% | 239 | 25.7% |
| *cis* and *trans* | 1,832 | 80.8% | 511 | 54.9% |
| opposite **CIS** *trans* | 116 | 5.1% | 127 | 13.6% |
| opposite *cis* **TRANS** | 93 | 4.1% | 49 | 5.3% |
| same **CIS** *trans* | 24 | 1.1% | 39 | 4.2% |
| same *cis* **TRANS** | 44 | 1.9% | 28 | 3.0% |
| compensatory | 1,555 | 68.6% | 268 | 28.8% |

## Discussion

In this study we investigated thermal tolerance and the underlying expression and regulatory networks to identify traits and mechanisms that may contribute to hybrid tiger salamander advantages in the wild. First, we compared the ability of CTS, BTS and hybrid individuals to tolerate acute thermal stress by assaying CTMax. This yielded significant differences in mean CTMax between the combined parental group and hybrids, but not between the two parental species. It also indicated an increased variance in hybrids compared to the parental group, which was similarly not detected between parent species. Critically, many hybrids were able to tolerate much greater heat stress before suffering physiological failure than either CTS or BTS. Second, we analyzed differential gene expression to identify differences in genomic response to temperature stress. Our results suggest substantial differences between the tissue-specific expression response of CTS and BTS to thermal stress, despite their equivalent CTMax values. These underlying differences, when combined in hybrids, may contribute to the ability of some hybrid individuals to tolerate extreme temperatures. Third, we scanned for patterns of allele specific expression that may explain how and why some hybrids exhibit these transgressive CTMax phenotypes, as well as the considerable variation in thermal tolerance among F1 hybrids. Finally we used evidence from both differential gene expression and allele specific expression analyses to evaluate the genomic mechanisms that drive these exaggerated hybrid traits.

### Physiological Response (CTMax)

We did not detect a difference in CTMax between CTS (33.77°C ± 0.39) and BTS (33.64°C ± 0.43). This equivalency may result from the physiological constraints that high temperatures impose (Huey & Bennett, 1987; Huey et al., 2012; W. I. Lutterschmidt, 1997; Markle, 2015; Youssef, Adolph, & Richmond, 2008). A recent meta-analysis concluded that critical thermal limits are often constrained compared to other physiological tolerances at a global scale (Sunday, Bates, & Dulvy, 2011). However, thermodynamic models have demonstrated that conserved phenotypes, such as thermal tolerance, may mask significant variation in their underlying genetic mechanisms (López-Maury, Marguerat, & Bähler, 2008). These mechanistic differences that result when species evolve in isolation may incorporate unique, roughly equivalent strategies for tolerating stressful temperatures. If such independent lineages come back into contact and hybridize, their unique mechanisms can combine to produce phenotypes that exceed either parental species (ex: Alter et al., 2017; Bidani et al., 2007; Perry et al., 2001).

Hybrid tiger salamanders have a greater mean thermal tolerance (35.24°C ± 0.53), than the combined parental group (33.69°C ± 0.292). Furthermore, we observed six hybrid individuals that maintained physical function at temperatures greater than the max observed in a roughly equal sample of CTS or BTS combined, suggesting that 40% of individuals exhibited transgressive thermal tolerance capacity. This 1.77°C mean increase in thermal tolerance of the six transgressive hybrids is biologically meaningful given the hot arid conditions that these salamanders endure in breeding ponds and during summer aestivation. This increased resistance may play a critical role in the differential survival of hybrid salamanders that has been well-documented in the wild, which may intensify given current projections of climate change in CTS habitat (Searcy & Shaffer, 2016).

### Differential Gene Expression

Differential expression (DE) analysis revealed substantial differences in gene expression between CTS and BTS in response to acute thermal stress. In muscle tissue we observed a greater expressional response to heat stress in CTS (359 genes) than in BTS (16 genes). The reverse is true in the brain tissue, where CTS had only 4 DE genes and BTS had 14. These data suggest differences in the number of genes that respond to thermal stress within each tissue. Additionally, we see an inverted pattern across tissue types, where CTS exhibits a greater expressional response in muscle, while BTS shows a greater response in the brain. Previous studies have also used patterns of gene expression to identify alternative mechanisms and pathways that species use to abate thermal stress. In nonmodel species, the majority of these experiments have focused on marine organisms (ex. Marine snail: Gleason and Burton, 2015; Fishes (review): Logan and Buckley, 2015; Abalone: Shiel et al., 2015; Trout: Tan et al., 2016; Salmon: Tomalty et al., 2015). These studies have uncovered many genes that are differentially expressed in response to heat stress, and genes that are uniquely expressed in species or populations with improved heat tolerance. Furthermore, differences in the number of genes that respond to thermal stress demonstrate differences in the underlying thermal tolerance mechanisms. Therefore, the differences we see in tissue specific gene expression between CTS and BTS support the interpretation that these two lineages have evolved different thermal abatement strategies.

### Transgression and Variation in Hybrids

Hybrid salamanders exhibited a greater range of thermal tolerance capabilities than the combined parental group. The variation in thermal tolerance of these primarily F1 hybrids underscores the complexity of mechanisms driving this apparent heterosis. Previous studies have demonstrated a wide range of genomic mechanisms that produce transgressive phenotypes, many of which conclude that a combination of processes are responsible (Comings & MacMurray, 2000; Pea, Ferron, Gianfranceschi, Krajewski, & Enrico Pè, 2008). In a review of human studies Comings and MacMurray (2000) concluded that the most likely mechanism of heterosis functions beyond the classical dominance and overdominance models, acting instead on the whole cell or whole organism. At this level many complex epigenetic factors may be influencing the transgressive patterns seen in hybrids, including differences in epigenetic methylation (Lauss et al., 2018) and expression of small regulatory RNAs (Groszmann et al., 2011; Shivaprasad, Dunn, Santos, Bassett, & Baulcombe, 2012). Since most of these mechanisms generate differences in gene expression at the whole gene and/or allele specific level, we analyzed allelic imbalance to better understand these transgressive and variable traits in F1 hybrids.

The F1 hybrids in our study exhibited relatively high levels of static ASE with 30.7% of genes in the muscle and 15.7% in the brain with biased expression. These values are high, though within the range of other studies (Keane et al., 2011). In addition to static ASE, we also examined ASE that changed in response to thermal stress. We found 3.1% of genes in the muscle and 0.4% in the brain that altered their expression of parental gene copies in response to thermal stress. This suggests a causal relationship between ASE and thermal tolerance, where increased bias in allelic expression may augment thermal tolerance in F1 individuals.

We further investigated the relationship between ASE and thermal tolerance by modeling CTMax as a function of each individual’s proportion of genes with significant ASE. We found strong support for this with non-condition dependent (static) ASE in the brain (Figure 4a), suggesting that temperature tolerance is influenced by ASE in this tissue. We also found some support for a correlation between temperature dependent ASE and CTMax in the muscle tissue (Figure 4b), although our limited sample size likely contributed to its marginal non-significance. This pattern supports the intriguing idea that thermal tolerance is affected by an individual’s degree of allelic expression imbalance. Furthermore, the tissue-specific differences in the correlation between CTMax and ASE again support the hypothesis that alternative mechanisms are employed by CTS and BTS in specific tissues to help mitigate thermal stress. It is also interesting to note that increased proportions of BTS-biased ASE in the brain correlates with improved thermal tolerance. Similarly, BTS exhibits the greatest overall expressional response to temperature in brain tissue, suggesting that BTS may have improved adaptations to thermal stress in this tissue. These interactions between allele specific expression and expression of phenotypic traits have been documented in other species as well, though the causal relationship remains poorly understood (Cotroneo et al., 2006; He et al., 2006; Keane et al., 2011). Together, these patterns of ASE suggest unbalanced regulation of parental gene copies in F1 hybrids. We therefore examine mechanisms that regulate gene expression to understand the abundance of ASE in hybrid salamanders.

### Genomic Mechanisms

Analysis of allele specific expression (ASE) revealed substantial differences in gene regulation between muscle and brain tissue. Comparing the level of expression of each gene in CTS and BTS with the two parental copies of that gene in F1 hybrids allows us to identify the regulatory mechanism that governs the expression of that gene. In the brain we observed fewer genes with only *trans-* (4.0%) than genes with only *cis-* (15.2%) regulatory elements. In contrast, we observed the inverse in muscle tissue, with 25.7% *trans-* only vs 19.4% *cis-* only derived differences in gene expression. This reciprocal pattern is similar to the tissue-specific patterns observed in overall expression with response to heat, again suggesting that differences in tissue-specific thermal abatement strategies may be fueling the patterns observed in hybrids. This pattern of tissue specific differences in gene regulation has been documented in other species as well (Barbeira et al., 2018). A large expression QTL study in humans found that regulatory mechanisms differed substantially by tissue within the same individual (Consortium, 2015). In a review, Wray (2007) suggested that tissue specific differences in gene regulation represent an essential evolutionary process that enables selection to act on specific tissues with different requirements. This suggests that different functional constraints imposed on brain and muscle tissue may offer insight into their observed regulatory differences. One explanation is that *trans-*acting mutations in the brain may lead to disastrous pleiotropic effects that are detrimental due to the brains conserved and specialized function (Duret & Mouchiroud, 2000). Therefore, regulatory evolution in the brain may favor *cis-* mutations which are localized and thus more heritable (Price et al., 2011). Conversely, these restrictions may be relaxed in muscle tissue where pleiotropic effects that result from *trans-* acting mutations may be less detrimental, or even beneficial. Support for this hypothesis in the BTS/CTS system may be seen in the relatively higher degree of compensatory mutations in the brain (68.6%) than in the muscle (28.8%). This suggests that gene regulation in the brain resists changes in overall expression (Landry et al., 2005; Mack, Campbell, & Nachman, 2016).

The enrichment of *trans-* regulatory elements in hybrid muscle tissue may reflect the evolutionary and demographic histories of CTS and BTS. It is often thought that *cis-* regulation should be predominant in the wild, since its heritable nature improves the efficiency of natural selection, allowing traits to evolve more rapidly in response to external pressures (Wray, 2007). However, organisms with small population sizes may experience a greater degree of genetic drift, reducing the efficiency of natural selection to promote the accumulation of *cis-* regulatory factors (Gilad, Oshlack, & Rifkin, 2006; Parsch & Ellegren, 2013). This is likely the case with CTS, which is restricted to several small populations in CA, many of which have extremely low effective population sizes (Shaffer and McCartney-Melstad, unpublished). In contrast, BTS has a large range spanning much of North America, extending from Northern Mexico to Southern Canada. A similar study on *Drosophila* found a greater fraction of *trans-* regulated genes likely driven by the restricted population size of one of the parental species (McManus et al., 2010).

Variation in hybrid expression and phenotype may result from extensive compensatory regulation. Compensatory regulation was detected at high levels in both the brain (68.6%) and muscle (28.8%) tissue. These are comprised of genes that exhibit significant ASE in hybrids, yet similar overall expression between CTS and BTS. Additionally, regulation where *cis-* and *trans*-were both detected but functioned in opposite directions (another form of incomplete compensation) was detected in both brain (9.2%) and muscle (18.9%). Both of these regulatory mechanisms can lead to expression patterns in hybrids that exceed the range of either parent. This phenomenon has been documented in many species including *Drosophila* (Michalak & Noor, 2003; Ranz, Namgyal, Gibson, & Hartl, 2004), *Arabidopsis* (Comai, Madlung, Josefsson, & Tyagi, 2003; Wang et al., 2004), and maize (Auger et al., 2005), all of which show transgressive gene expression patterns in hybrids. This process is thought to occur when mutations in *trans-* acting elements have a net benefit to an organism, but consequently have a detrimental, pleiotropic effect through the expression of other co-regulated genes. These negative effects are then reduced through subsequent mutations (likely *cis-*) that shift the expression of these other genes back to the original optimum through purifying selection. This mechanism maintains the benefit from the initial *trans-* mutation, while reducing the negative downstream effects it has on other genes. This may explain why genes exhibit equivalent levels of expression in CTS and BTS, yet have different values in the F1 hybrids. The decoupling of these compensatory mutations result in novel phenotypes, which may explain the variability in hybrid CTMax, and the ability of some hybrids to handle greater heat stress than either parent. This process is akin to transgressive segregation, however here, recombination in F2 individuals is not required to produce novel phenotypes (Rieseberg et al., 1999). Rather, compensatory evolution has occurred at the expression level, and the *trans-* acting factors that influence expression are not equally inherited by F1 offspring, producing a variable phenotype similar to the recombinant effect of meiosis. The heritability of these F1 expression patterns is still largely unknown, and requires additional study. However we do see that two of the three backcross/F2 heat-stressed individuals included in the CTMax experiment also exhibit transgressive thermal tolerance, suggesting that this is not purely a phenomenon limited to first-generation hybrids. Furthermore, two human expression studies found that 29% of surveyed genes have heritable genetic components (Schadt et al., 2003), and that 15% of the variation in gene expression is heritable across multiple tissue types (Price et al., 2011). These studies suggest that there is significant heritability in gene expression. If the expression-level traits that we document here are heritable, then selection on increased CTMax may lead to hybrid salamanders with enhanced thermal tolerances in the wild. Future work should examine the CTMax of wild hybrid salamanders that have undergone multiple generations of selection, to explore whether this pattern of increased thermal tolerance, or at least variation in CTMax, exists in the wild.

### Conclusion

Our study demonstrates how an apparently conserved phenotype shared between two species may conceal significant differences that have accumulated since the species diverged from a common ancestor. Our approach leverages the power of next generation transcriptome sequencing to uncover the genomic mechanisms that contribute to cryptic differences between species. New genomic tools, combined with the increased availability of reference genomes for even the most recalcitrant non-model systems, like salamanders, are enabling analyses into complex regulatory mechanisms in species that can be of great conservation concern.

We have identified thermal tolerance as a potential factor influencing hybrid tiger salamander success. We found substantial variation in hybrid CTMax, with many individuals tolerating hotter temperatures than either parental species, and differences in gene expression between CTS, BTS and hybrids suggesting mechanistic differences in response to thermal stress. This is corroborated by biased allele specific expression in hybrid individuals, and is consistent with the interpretation that developmental-system drift has occurred between CTS and BTS (True & Haag, 2001). These differences appear to confer an increase in thermal tolerance to some of the resulting hybrids that may be a key to their previously documented increased fitness (Fitzpatrick et al., 2009).

These results suggest that hybrids may be more fit than native CTS in response to future climate change. This may have implications for management, depending on the viability of native CTS in the face of warming temperatures. If native CTS populations decline due to temperature stress, then hybrids may become the only viable option for salamander persistence in the hotter regions of the species’ range. Previous experimental work has demonstrated that although hybrids have a disruptive effect on experimental vernal pool ecosystems, that disruption is far greater when no *Ambystoma* larvae are present (Searcy et al., 2016). Given this, it may be reasonable to protect hybrids, while simultaneously attempting to restore breeding pool environments to more natural, vernal-pool conditions (e.g. Fitzpatrick & Shaffer, 2007b; Wayne & Shaffer, 2016). This strategy may allow for natural selection for other, predominantly native, traits, effectively reducing the non-native alleles to just those that confer adaptive temperature tolerance. This is especially relevant given recent predictions of large shifts in temperature-dependent habitat suitability by 2070 throughout the CTS range (Searcy & Shaffer, 2016). One of the strong conclusions from this study is the prediction that the only suitable habitat for CTS will be in the central coast region where the current hybrid zone is expanding (Searcy & Shaffer, 2016), suggesting that hybrid populations may occupy the only viable habitat for CTS given persistent climate change. More work needs to be conducted on the current hybrids to determine if increased thermal tolerance is indeed evolving in the hybrid zone.

Future studies should target naturally occurring hybrids to identify CTMax values and the resulting gene expression at this critical temperature. By comparing the data we have presented here to wild hybrid populations rather than just F1s, we can test the hypothesis that selection favors hybrids with higher thermal tolerances in the wild, and investigate the underlying gene expression patterns facilitating this adaptation. Higher CTMax values and large expression differences would be consistent with the prediction that hybrid regulatory mismatch produces some individuals with enhanced thermal tolerance, which affords them greater fitness in the wild. In addition, physiological models that predict population extinction probability using climate change projections can be used to estimate the effect these differences in thermal tolerance may have on population viability. It is conceivable that higher thermal tolerance conferred via hybridization may shield CTS-hybrid populations from future climate change which would otherwise push CTS beyond its thermal limit. If so, it may be time to consider full protection for thermally-tolerant hybrids as the best option to retain ecologically similar, but not identical, CTS on their remaining natural landscapes.

## Acknowledgements

We thank Drs. C. Searcy, R. Voss, J. Smith for insight into methods and analyses. We thank the Shaffer and Grether Lab for insightful comments. We thank E. Toffelmier for tremendous assistance in the lab and with the manuscript, and the La Kretz Center for California Conservation and the Ecology and Evolutionary Biology Department at UCLA, the Natural Communities Coalition, and the University of California Conservation Genomics Consortium (CA-16-376437) for partial funding.

## Data Accessibility

All trimmed RNA sequences along with generated count and meta data have been deposited in the NCBI’s Gene Expression Omnibus (Edgar, Domrachev, & Lash, 2002) and are accessible through GEO Series accession number GSE137607 (https://www.ncbi.nlm.nih.gov/geo/query/acc.cgi?acc=GSE137607).

## Author Contributions

R.D. Cooper: Designed and performed experiment, analyzed data and wrote the manuscript. H.B Shaffer: Contributed to the project design and analysis, and manuscript editing.

## Conflict of Interest

We have no conflict of interest to report

